## Supplementary figures and tables for "TWEAK/Fn14 signalling driven super-enhancer reprogramming promotes pro-metastatic metabolic rewiring in Triple-Negative Breast Cancer"

### Supplementary material

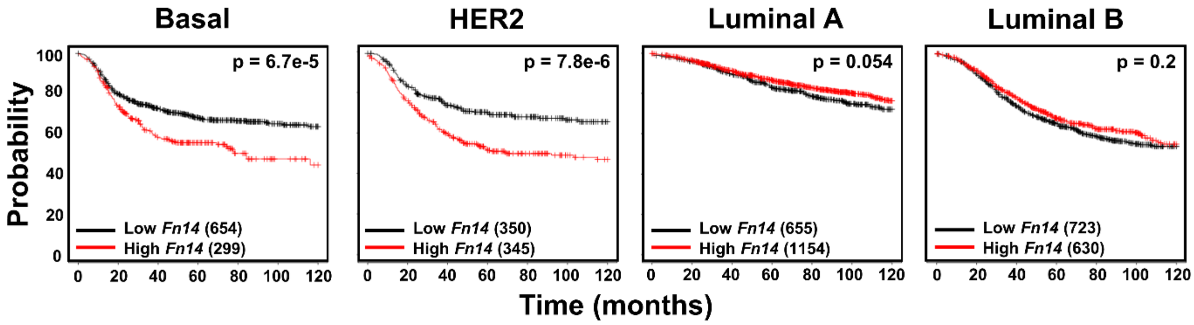


**Supplementary Figure 1. High Fn14 expression confers worse survival in Basal-like and HER2 patients.** Kaplan Meier plot depicting the relapse-free survival of *Fn14* high and low Basal, HER2, Luminal A and Luminal B breast cancer patients.

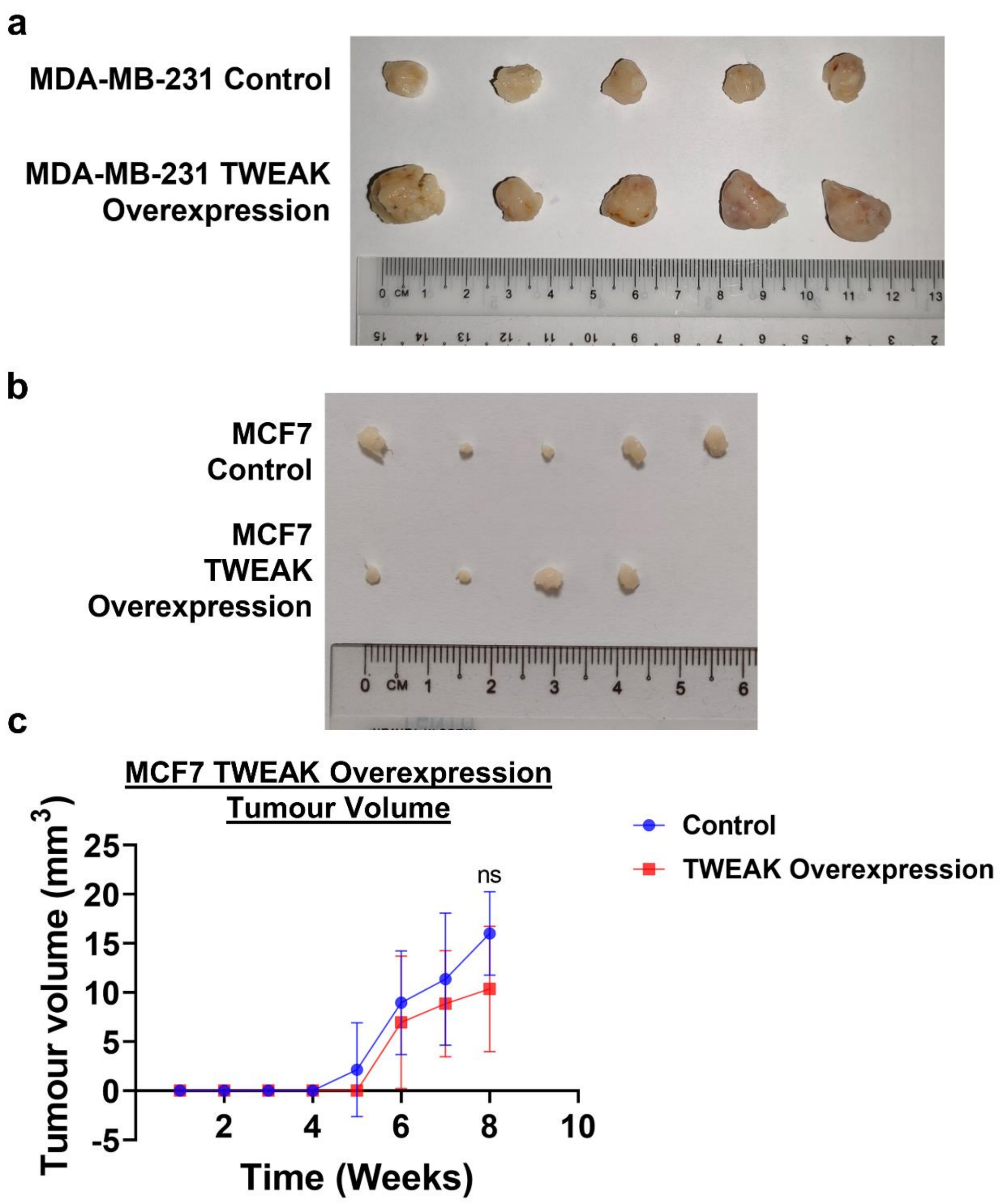


**Supplementary Figure 2. Persistent Fn14 activation promotes tumour growth in TNBC but not ER-positive breast cancer cells.** Images of tumours extracted from mice 8 weeks after being injected with (a) luciferase expressing MDA-MB-231 cells (n=5 biological replicates) and (b) MCF7 cells (n=5 biological replicates) harbouring the overexpression control or TWEAK overexpression. There was no tumour growth in the 5th MCF7 TWEAK Overexpression mouse. (c) Tumour growth plot depicts the weekly average tumour volume (mean ± s.d) from mice injected with MCF7 cells overexpressing luciferase and overexpression control or TWEAK. T-test was used for statistical analysis.

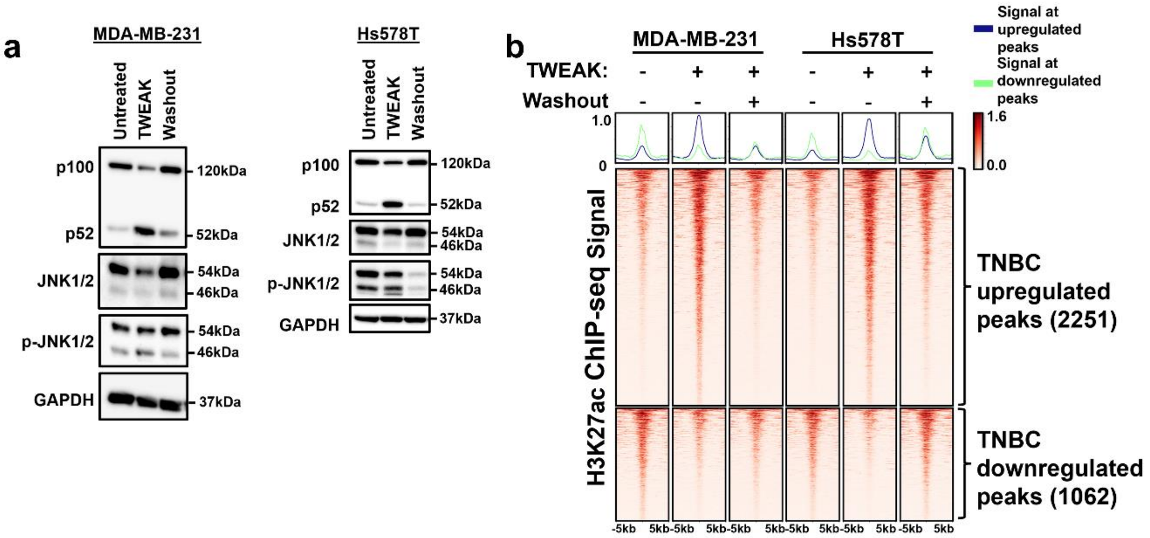


**Supplementary Figure 3. Wash off ameliorates TWEAK-driven non-canonical NF-κB pathway, JNK pathway and enhancer activation in TNBC.** (a) NF-κB and JNK pathway signalling regulators analysed by western blotting in untreated, TWEAK-treated and TWEAK-treated with 96h wash-off MDA-MB-231 and Hs578T cells. (b) H3K27ac ChIP-seq signals of untreated, TWEAK-treated and TWEAK-treated with 96h wash-off MDA-MB-231 and Hs578T cells at TWEAK/Fn14-driven differential TNBC enhancer sites.

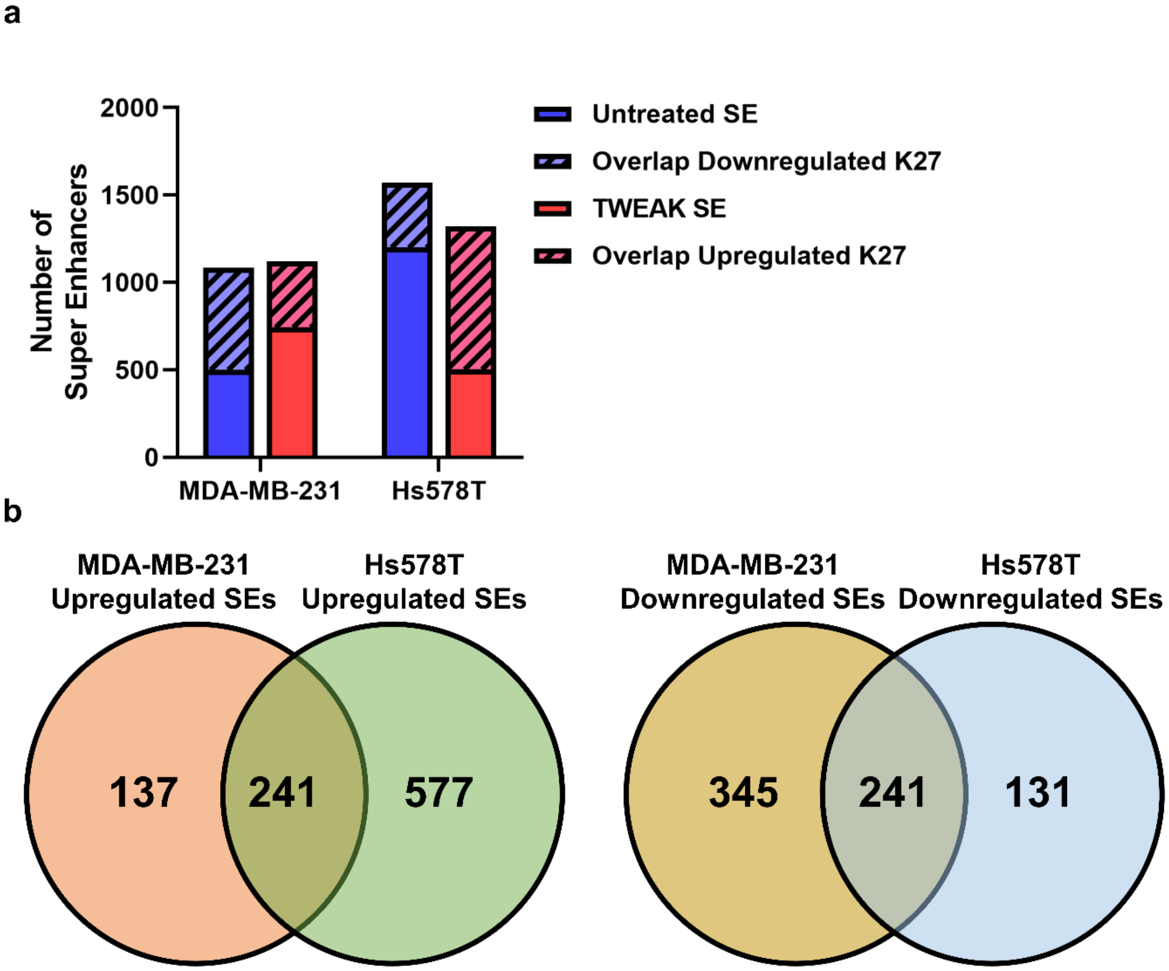

**Supplementary Figure 4. TWEAK/Fn14 signalling is highly involved in TNBC SE regulation.** (a) Bar plot depicting the number of SEs called in MDA-MB-231 and Hs578T cells with and without TWEAK treatment, as well as the number of SEs that overlap up- and down-regulated TNBC H3K27ac ChIP-seq peaks. (b) Venn diagrams depicting the number of common and cell line specific TWEAK/Fn14 regulated SEs between MDA-MB-231 and Hs578T cells.

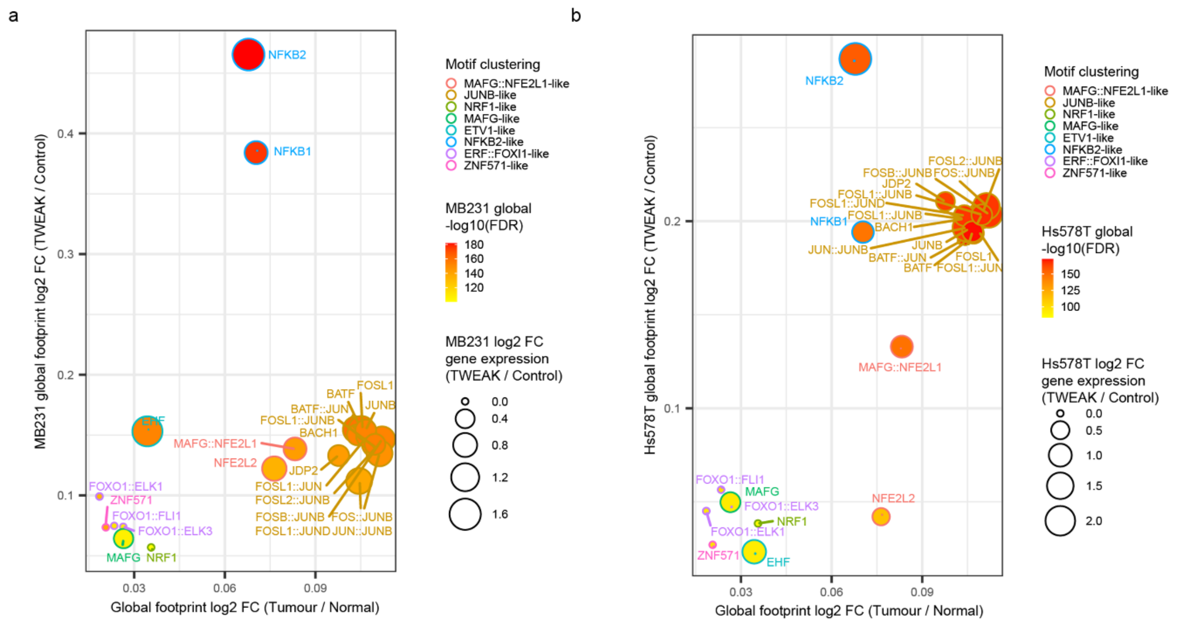

**Supplementary Figure 5. Global TF DNA binding dynamics in TWEAK stimulated MDA-MB-231, Hs578T and Fn14-high TNBC patient tumours.** (a) MDA-MB-231 and (b) Hs578T global log2 fold changes for footprints and gene expression are plotted with Fn14 high TNBC patient tumour global footprint log2 fold changes. TF binding changes exhibiting p-values in the upper half of the distribution are selected and filtered for consistent global footprint and gene expression dynamics across cell lines and tumours. Motifs are clustered based on similarity via TOBIAS.

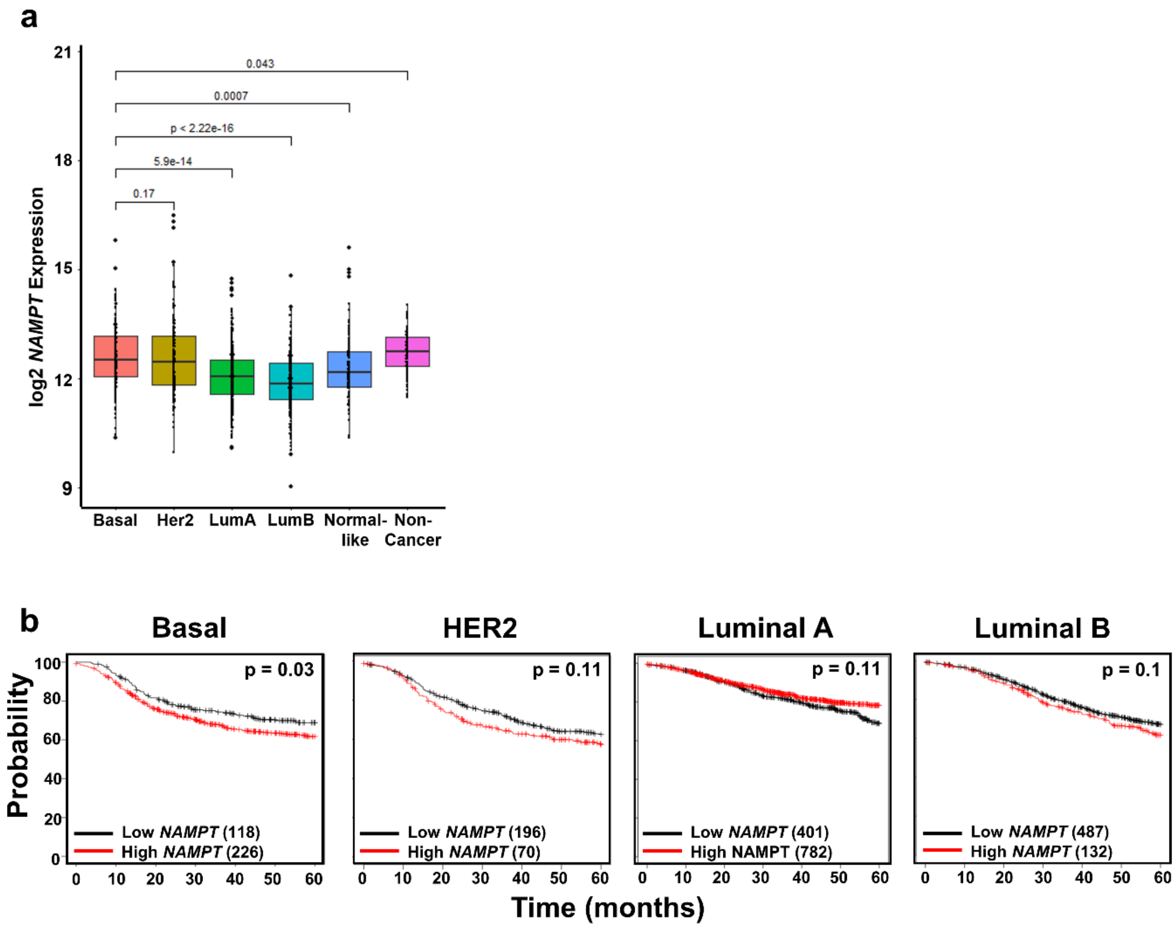


**Supplementary Figure 6. NAMPT is overexpressed in ER-negative breast tumours and its overexpression confers worse survival selectively in basal-like breast cancer patients.** (a) Relative *NAMPT* gene expression levels in Basal-like, HER2, Luminal A, Luminal B, Normal-like and non-cancer patient samples from the TCGA BRCA RNA-seq dataset. (b) Kaplan Meier plot depicting the relapse-free survival of *NAMPT* high and low Basal-like, HER2, Luminal A and Luminal B breast cancer patients.

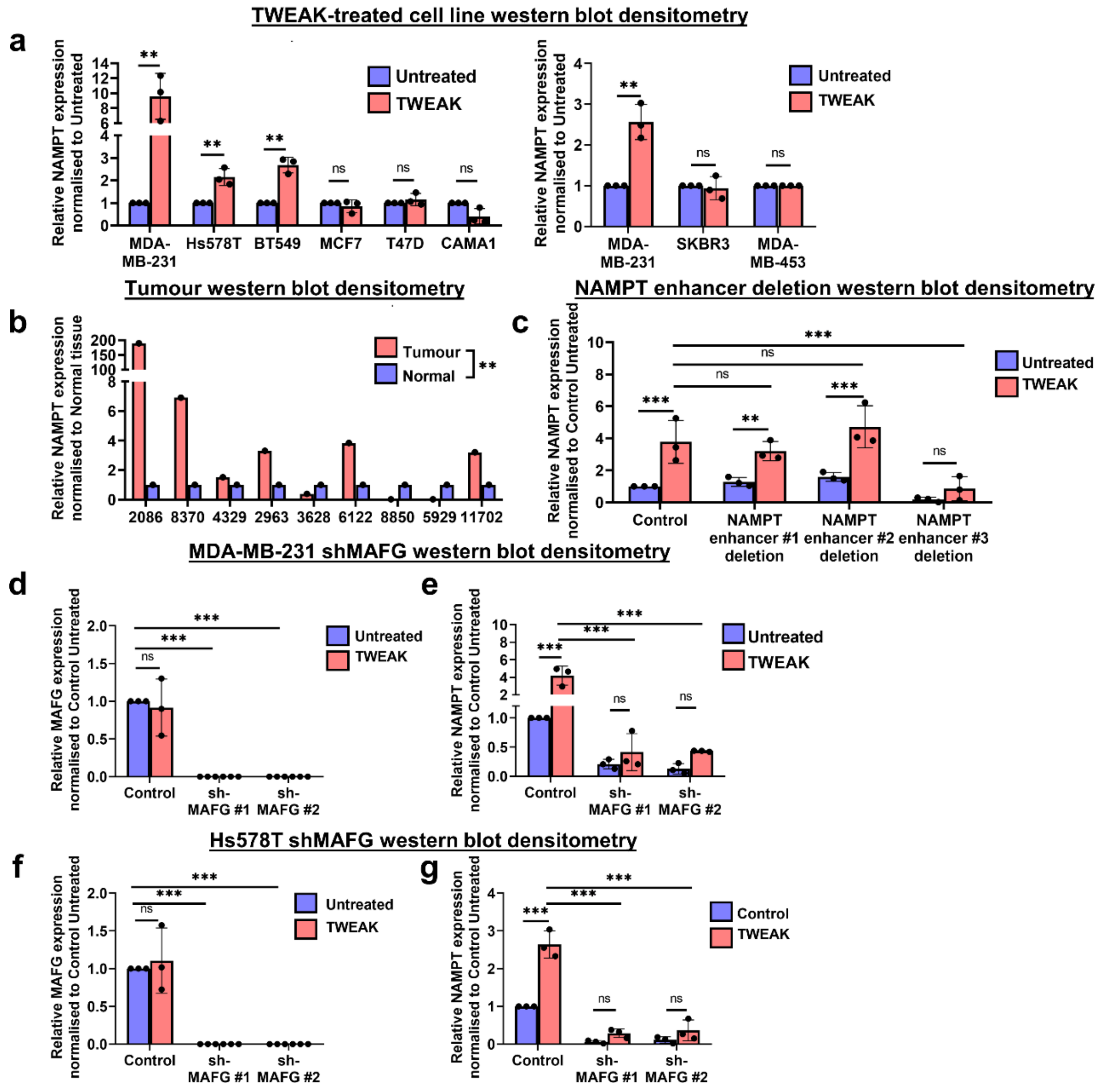


**Supplementary Figure 7. Western blot densitometries.** Relative NAMPT expression to GAPDH, (a) normalised to each cell line's untreated sample in Figure. 6a, (b) normalised to each tumour sample's matched normal tissue in Figure 6b and (c) normalised to the control untreated sample. Relative (d) MAFG and (e) NAMPT expression to GAPDH, normalised to the control untreated MDA-MB-231 sample in Extended Data Figure 8. Relative (f) MAFG and (g) NAMPT expression to GAPDH, normalised to the control untreated Hs578T sample in Extended Data Figure 8. T-test was used for in Tumour vs Normal statistical analysis while Two-way ANOVA was used for statistical analysis in all other plots, where *P < 0.05; **P < 0.01; ***P < 0.001.

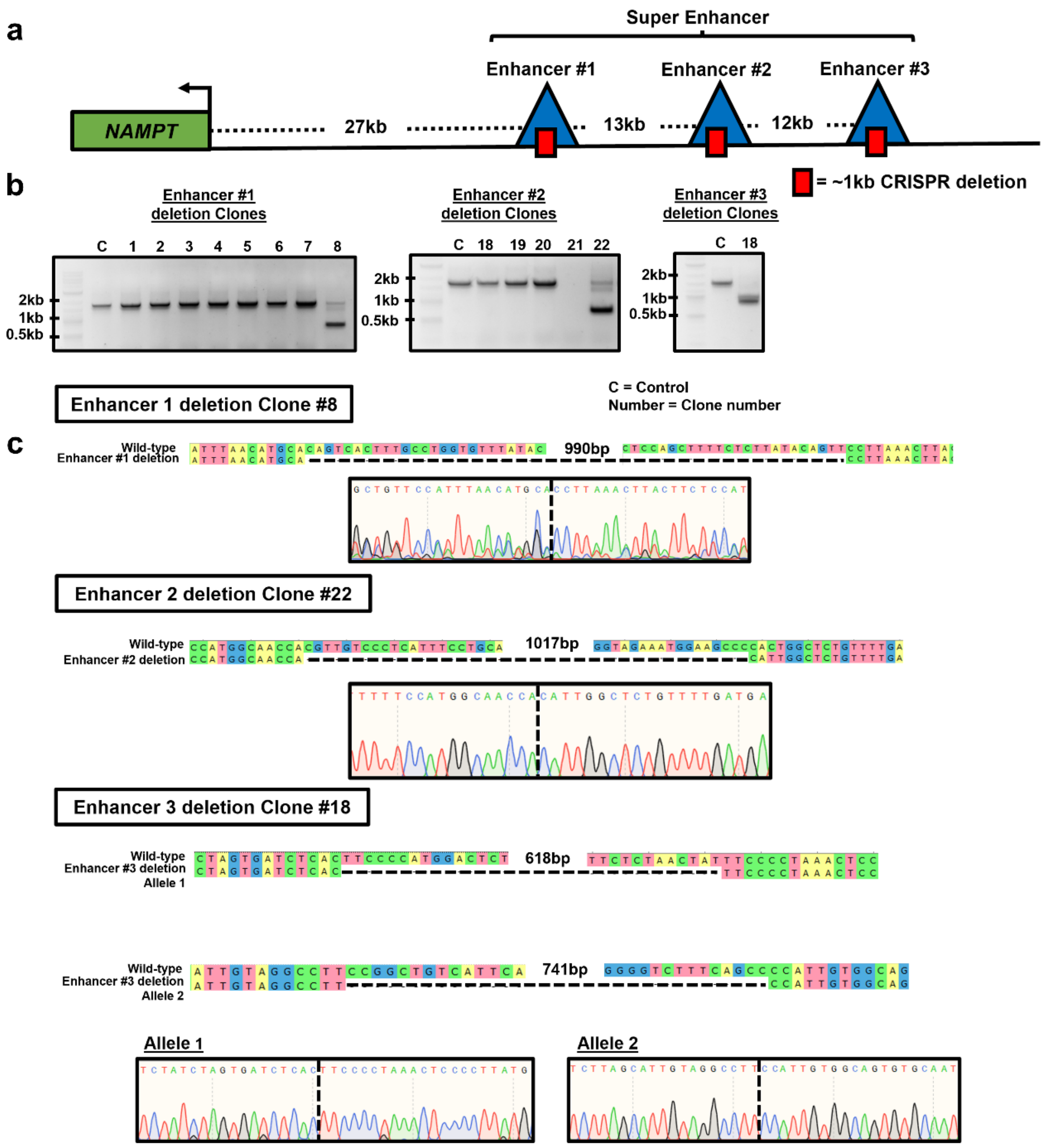


**Supplementary Figure 8. *NAMPT* enhancer CRISPR deletion strategy and genotyping of CRISPR-edited clones.** (a) Diagram depicting the *NAMPT* locus and the locations where the 1kb CRISPR/Cas9 deletions were made. (b) Gel electrophoresis images of MDA-MB-231 clones harbouring successful deletions in *NAMPT* enhancer #1, #2 and #3. (c) Sanger sequencing tracks showing the regions in *NAMPT* enhancer #1, #2 and #3 that were successfully deleted in MDA-MB-231 clones.

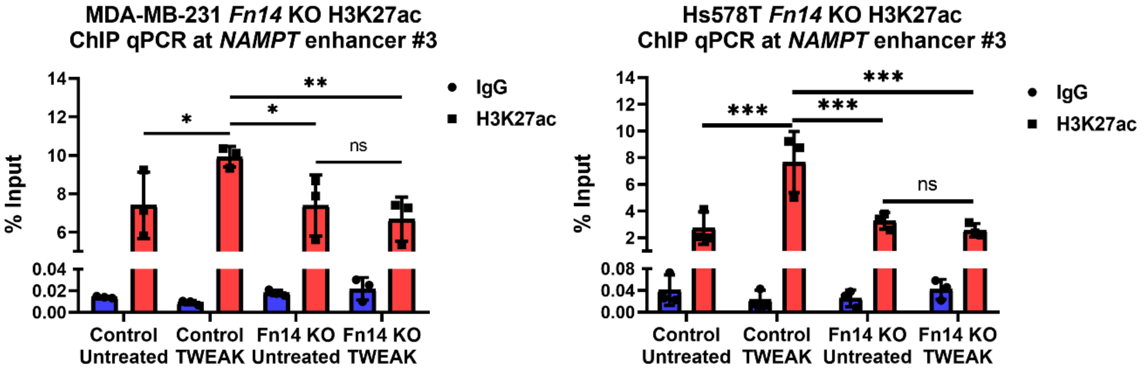


**Supplementary Figure 9. Fn14 KO abolishes TWEAK/Fn14-driven *NAMPT* enhancer #3 regulation.** Plot depicting the IgG and H3K27ac ChIP qPCR %input at *NAMPT* enhancer #3 in control and Fn14 KO MDA-MB-231 and Hs578T cells with and without TWEAK treatment (mean ± s.d). Data shown represent n=3 biological replicates. Two-way ANOVA was used for statistical analysis where *P < 0.05; **P < 0.01; ***P < 0.001.

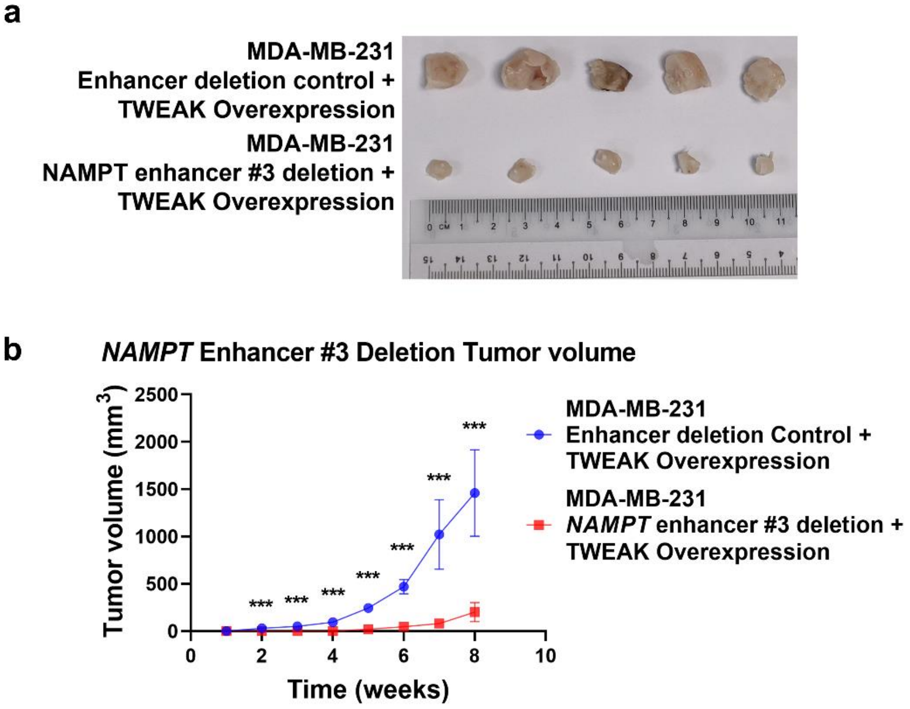


**Supplementary Figure 10. *NAMPT* enhancer #3 deletion ameliorates TWEAK/Fn14-driven TNBC tumour growth.** (a) Images of tumours extracted from mice 8 weeks after being injected with luciferase and TWEAK overexpressing MDA-MB-231 cells that either harbour the *NAMPT* enhancer #3 deletion or without (n=5 biological replicates). (b) Tumour growth plot depicts the weekly average tumour volume (mean ± s.d) from the mice. Performed in n=5 biological replicates. T-test was used for statistical analysis where *P < 0.05; **P < 0.01; ***P < 0.001.

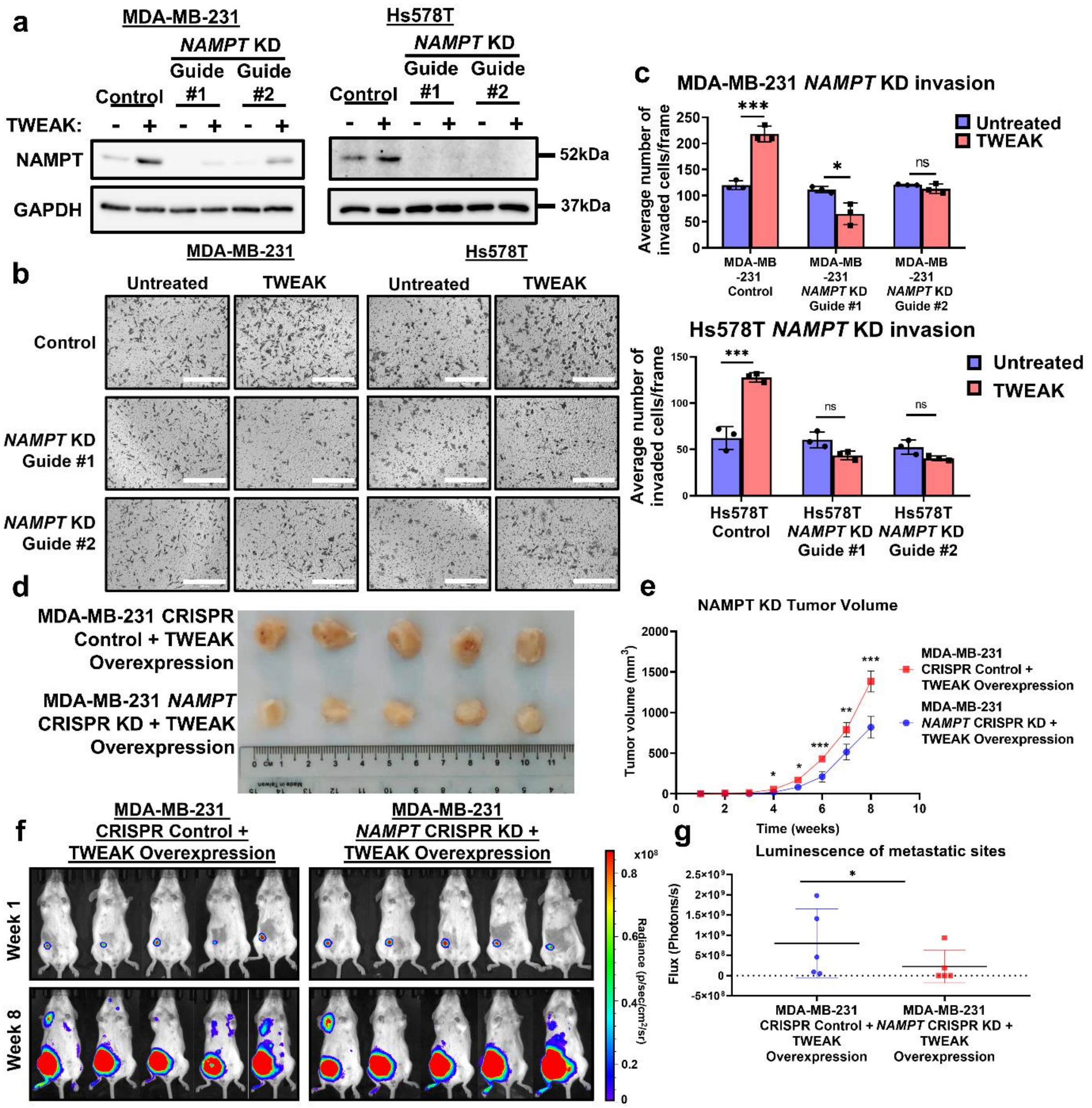


**Supplementary Figure 11. NAMPT is a critical regulator of TWEAK/Fn14-driven tumour growth and metastasis in TNBC.** (a) NAMPT expression analysed by western blotting in control and *NAMPT* KD MDA-MB-231 and Hs578T cells, with and without TWEAK treatment. (b) Transwell invasion assay was performed in control and *NAMPT* KD MDA-MB-231 and Hs578T cells, with and without TWEAK treatment. Representative images from n=3 biological replicates are shown. Scale bars: 400μm. (c) Representative plot depicts the average number of invaded cells/frame (mean ± s.d) from n=3 biological replicates, across four fields per replicate. (d) Images of tumours extracted from mice 8 weeks after being injected with luciferase and TWEAK overexpressing control and *NAMPT* KD MDA-MB-231 cells (n=5 biological replicates). (e) Tumour growth plot depicts the weekly average tumour volume (mean ± s.d). Performed in n=5 biological replicates. (f) IVIS tracking of mice injected with luciferase and TWEAK overexpressing control and *NAMPT* KD MDA-MB-231 cells. Representative bioluminescent images of the animals were taken at 1 and 8 weeks after orthotopic xenograft. (g) Plot depicts total flux at the metastatic sites of each animal after 8 weeks in n=5 biological replicates. Two-way ANOVA was used for statistical analysis in *in vitro*

invasion assay. T-test was used for statistical analysis in *in vivo* assays. *P < 0.05; **P < 0.01; ***P < 0.001.

| Name | Sequence |
|---|---|
| shMAFG #1 | CCAGCGTCATCACAATAGTAA |
| shMAFG #2 | CCTCAGAGAACGCCAGCATGA |

**Supplementary Table 1: shRNA sequences used.**

| Name | Sequence |
|---|---|
| NAMPT enhancer #1 F | ACCGCTGTTTATTGGGCATTT |
| NAMPT enhancer #1 R | CCCACTCCACTGCTTGCTAG |
| NAMPT enhancer #2 F | CCCCACTGGCTCTGTTTTGA |
| NAMPT enhancer #2 R | CACTCAACCCCTTCTCCCAC |
| NAMPT enhancer #3 F | AGTGCTTCCCTTCCCTGAGA |
| NAMPT enhancer #3 R | GCGGAAGTAAGCATCGTGAC |

**Supplementary Table 2: ChIP qPCR primers.**
